# G6PD deficiency sensitizes metastasizing melanoma cells to oxidative stress and glutaminolysis

**DOI:** 10.1101/2021.11.11.468286

**Authors:** Arin B. Aurora, Vishal Khivansara, Ashley Leach, Jennifer G. Gill, Misty Martin-Sandoval, Chendong Yang, Stacy Y. Kasitinon, Divya Bezwada, Alpaslan Tasdogan, Wen Gu, Thomas Mathews, Zhiyu Zhao, Ralph J. DeBerardinis, Sean J. Morrison

## Abstract

The pentose phosphate pathway is a major source of NADPH for oxidative stress resistance in cancer cells but there is limited insight into its role in metastasis, when some cancer cells experience high levels of oxidative stress. To test this, we mutated the substrate binding site of Glucose-6-phosphate dehydrogenase (G6PD), which catalyzes the first step of the pentose phosphate pathway, in patient-derived melanomas. *G6PD* mutant melanomas had significantly decreased G6PD enzymatic activity and depletion of intermediates in the oxidative branch of the pentose phosphate pathway. Reduced G6PD function had little effect on the formation of primary subcutaneous tumors but when these tumors spontaneously metastasized the frequency of circulating melanoma cells in the blood and metastatic disease burden were significantly reduced. *G6PD* mutant melanomas exhibited increased levels of reactive oxygen species (ROS), decreased NADPH levels, and depleted glutathione as compared to control melanomas. *G6PD* mutant melanomas compensated for this increase in oxidative stress by increasing the production of NADPH through glutaminolysis. This generated a new metabolic vulnerability as *G6PD* mutant melanomas were more dependent upon glutamine as compared to control melanomas. The oxidative pentose phosphate pathway and compensatory glutaminolysis thus confer layered protection against oxidative stress during metastasis.

**Significance:** Melanoma metastasis is limited by oxidative stress. Cells that enter the blood experience high levels of ROS and usually die of ferroptosis. We found that melanoma cells become more dependent upon the oxidative branch of the pentose phosphate pathway to manage oxidative stress during metastasis. When pentose phosphate pathway function was disabled by *G6PD* mutation, the melanoma cells increased their utilization of malic enzyme, fueled by increased consumption of glutamine in the tricarboxylic acid cycle. Melanoma cells thus have redundant and layered protection against oxidative stress.

## Introduction

The pentose phosphate pathway is an important source of NADPH for oxidative stress resistance (1–5). The oxidative branch of the pentose phosphate pathway contains two enzymes that generate NADPH from NADP^+^, glucose 6-phosphate dehydrogenase (G6PD) and 6-phosphogluconate dehydrogenase (PGD) (Fig. S1). NADPH is an important source of reducing equivalents for oxidative stress resistance because it is used by cells to convert oxidized glutathione (GSSG) to glutathione (GSH), an abundant redox buffer. Complete deficiency for G6PD is embryonic lethal in mice (2, 6, 7) but hypomorphic G6PD mutations are common in certain human populations, perhaps because they protect against malaria (8, 9). These partial loss-of-function G6PD mutations are well tolerated in adults, though they sensitize red blood cells to hemolysis from oxidative stress under certain circumstances (10).

Several studies have reported a lower incidence and mortality for certain cancers in people with hypomorphic mutations in G6PD (11–14), suggesting that cancer cells depend upon G6PD to manage oxidative stress. Cells experience high levels of oxidative stress during certain phases of cancer development and progression, including during metastasis (15–17).

Antioxidant mechanisms thus promote the survival of cells during oncogenic transformation (18, 19) as well as during metastasis (15, 16). For example, relative to primary cutaneous tumors, metastasizing melanoma cells exhibit increased dependence upon the folate pathway (15), Monocarboxylate Transporter-1 (MCT1) (20), and glutathione peroxidase-4 (GPX4) (21), each of which directly or indirectly attenuate oxidative stress. By better understanding the mechanisms that confer oxidative stress resistance in cancer cells it may be possible to develop pro-oxidant therapies that inhibit cancer progression by exacerbating the oxidative stress experienced by cancer cells.

G6PD (22) or PGD deficiency (23–25) reduce the growth of some cancers, including melanoma, but G6PD deficiency has little effect on primary tumor formation by K-Ras-driven epithelial cancers (26). This is at least partly because loss of G6PD in these cancers leads to compensatory increases in the function of other NADPH-generating enzymes, including malic enzyme and isocitrate dehydrogenase (1). Nonetheless, pentose phosphate pathway function may increase during metastasis (20, 27–29) and higher G6PD expression is associated with worse outcomes in several cancers (30–32), raising the question of whether metastasizing cells are particularly dependent upon G6PD. G6PD is not essential for metastasis in a breast cancer cell line but it reduces their capacity to form metastatic tumors (26).

Melanoma cells exhibit little evidence of oxidative stress in established primary tumors but exhibit increased levels of ROS and dependence upon antioxidant mechanisms during metastasis (15, 20, 21). To test whether these cells also exhibit increased dependence upon the pentose phosphate pathway during metastasis we generated three G6PD mutant melanomas, including two patient-derived xenograft melanomas and one human melanoma cell line. G6PD deficiency had little effect on the formation or growth of primary subcutaneous tumors but significantly increased ROS levels and reduced spontaneous metastasis. G6PD deficient melanomas compensated by increasing the production of NADPH by malic enzyme. Melanoma cells thus have redundant layers of protection against oxidative stress during metastasis including the abilities to alter fuel consumption and antioxidant pathway utilization.

## Results

### Pentose phosphate pathway metabolites are enriched in metastases

To compare the levels of pentose phosphate pathway metabolites (Fig. S1) in subcutaneous tumors and metastatic nodules, we subcutaneously transplanted efficiently metastasizing melanomas obtained from three patients (M405, M481, and UT10) into NOD/SCID IL2Rγ^null^ (NSG) mice, allowing them to form primary subcutaneous tumors and to spontaneously metastasize. All of the melanomas were tagged with constitutive DsRed and luciferase, unambiguously distinguishing these cells from recipient mouse cells (Fig. S2A and S2B for flow cytometry gating strategies) and enabling the quantitation of metastatic disease burden by bioluminescence imaging (Fig. S2C-D). We dissected subcutaneous tumors and small metastatic nodules (<3mm in diameter) from the same mice and compared G6PD and PGD expression. G6PD and PGD were consistently expressed by both primary subcutaneous tumors and metastatic nodules at the RNA (Fig. 1A-B) and protein levels (Fig. 1C). Expression levels varied among tumors but did not differ in any consistent way between subcutaneous tumors and metastatic nodules.

**Figure 1:**
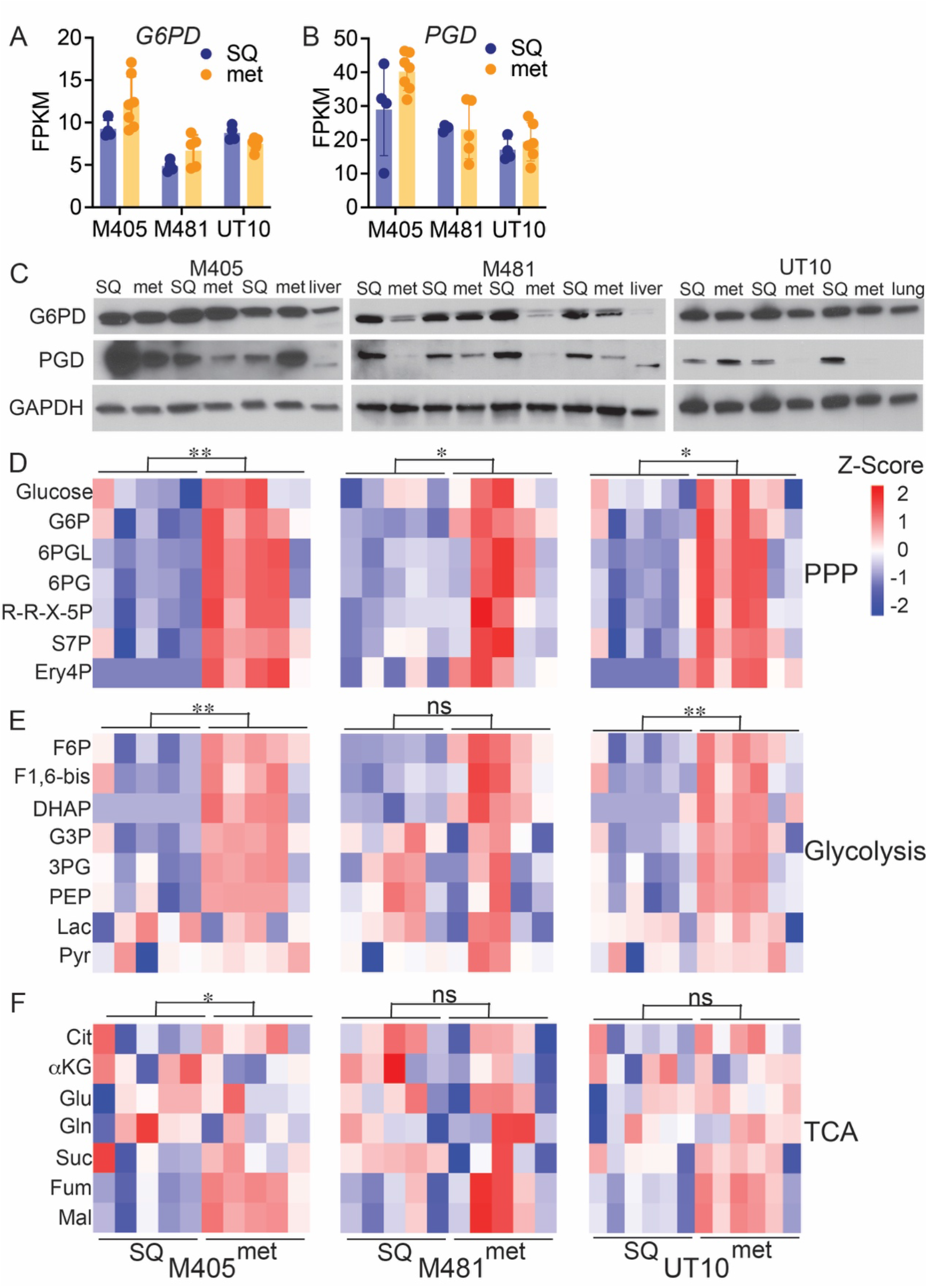
Pentose phosphate pathway metabolites are more abundant in metastatic as compared to subcutaneous primary melanomas. (A-C) Subcutaneous tumors (SQ) and metastatic nodules (met) were harvested from the same mice xenografted subcutaneously with M405, M481, and UT10 patient-derived melanomas. *G6PD* (A) and *PGD* (B) transcripts were assessed by RNAseq analysis (n=4-7 samples per melanoma). (C) Western blot analysis of G6PD and PGD in subcutaneous and metastatic tumors from mice xenografted with M405, M481, and UT10 melanomas. Each pair of adjacent subcutaneous and metastatic samples were obtained from the same mouse (n=3-4 mice/melanoma). GAPDH is shown as a loading control. Lysates from normal mouse liver or lung are included as controls. (D-F) Relative levels of pentose phosphate pathway (PPP), (D), glycolytic (E), and TCA cycle (F) metabolites in subcutaneous and metastatic tumor specimens obtained from mice xenografted with M405 (left), M481 (center), or UT10 (right) melanomas. Each column represents a different tumor from a different mouse. The statistical significance of each pathway was assessed by comparing the major effect between subcutaneous and metastatic tumors using a repeated measure two-way ANOVA. Data represent mean ±s.d. Statistical significance was assessed using Student’s t-tests (A-B), or a 2-way repeated measures ANOVA on the log2-transformed data (D-F). Statistical tests were two-sided. Multiple comparisons were adjusted by the FDR method. *P<0.05, **P<0.01, ns= not significant.

We performed a targeted metabolomic analysis on primary subcutaneous tumors and metastatic nodules and found that levels of pentose phosphate pathway metabolites were significantly higher in metastatic nodules as compared to subcutaneous tumors in all three melanomas (Fig. 1D). The levels of glycolytic intermediates were also significantly higher in metastatic nodules as compared to subcutaneous tumors in two of the melanomas (Fig. 1E). TCA cycle intermediates were elevated in metastatic nodules from one melanoma (Fig. 1F). The consistently higher levels of pentose phosphate pathway metabolites in most metastatic nodules raised the possibility that the pathway was more active in metastatic as compared to primary subcutaneous tumors.

### G6PD mutant melanomas

To test whether melanoma metastasis depended on the oxidative pentose phosphate pathway, we used CRISPR to make small deletions in exon 6 of *G6PD* in three melanomas including the A375 human melanoma cell line and two patient-derived melanomas (M481 and M214). Exon 6 encodes the substrate binding domain and mutations in exon 6 severely reduce G6PD enzymatic activity (33). We obtained three independently targeted clones for each melanoma and confirmed by sequencing that they had 66-68 base pair deletions in exon 6 (Fig. 2A). The *G6PD* mutant clones had greatly reduced levels of G6PD enzymatic activity (Fig. 2B).

**Figure 2:**
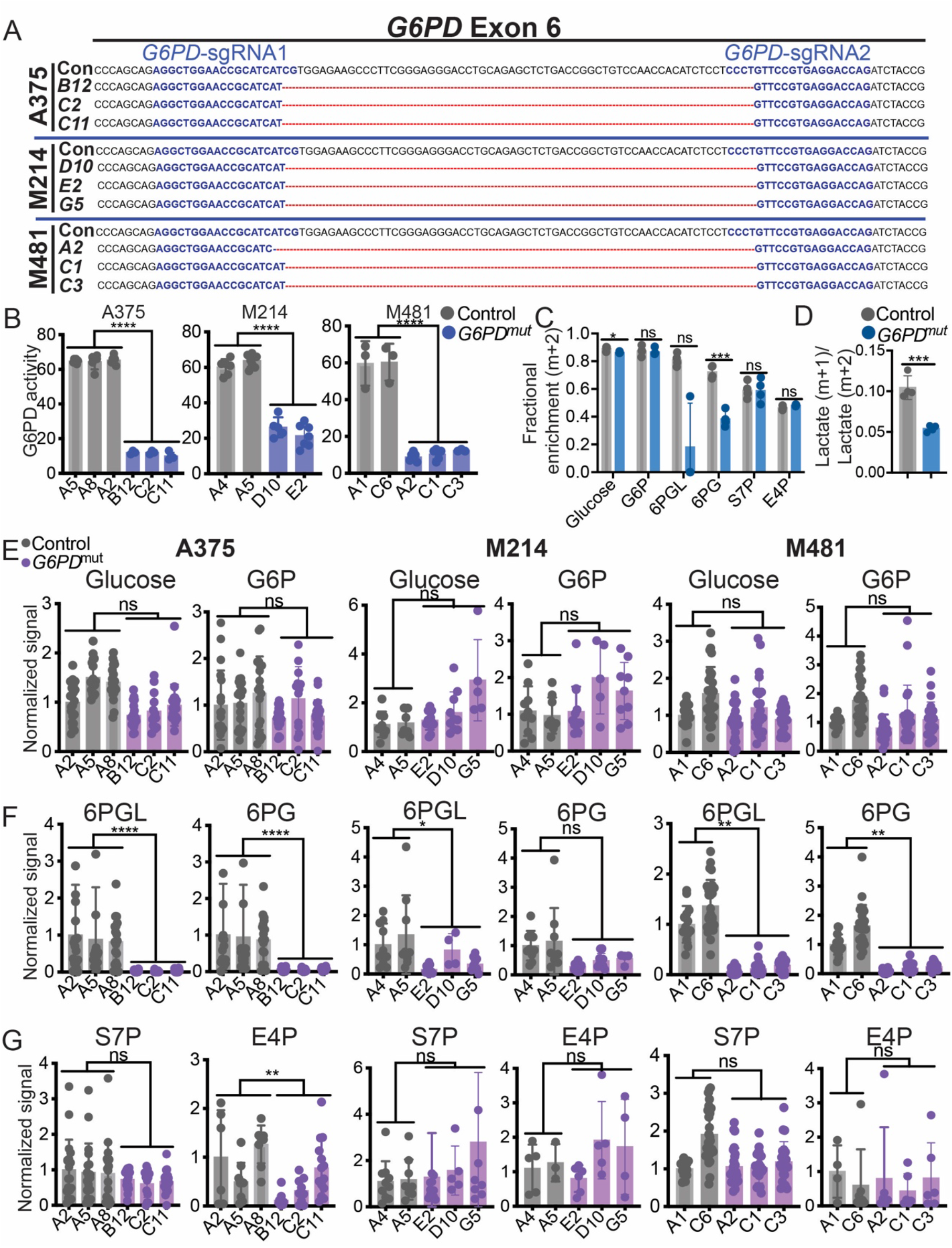
Generation of *G6PD* mutant melanoma cells with impaired oxidative pentose phosphate pathway function. (A) CRISPR editing of *G6PD* in three melanomas (A375, M214, and M481) to generate three independently targeted clones per melanoma with 66-68 base pair deletions in exon 6. A375 was derived from a female patient while M214 and M481 are from males. Guide sequences are highlighted in blue. The 66-68bp region deleted in the mutants is depicted by red lines. The control (Con) sequence is shown for reference. (B) G6PD enzymatic activity in subcutaneous tumors formed by *G6PD* mutant or control melanoma cells. (C-D) Tracing of [1,2-^13^C] glucose in *G6PD* mutant or control A375 melanoma cells to assess the activity of the pentose phosphate pathway. The data show the fractional enrichments (m+2) in glucose, G6P, 6PGL, 6PG, E4P, S7P (*C*) and the ratio of m+1 lactate to m+2 lactate (*D*) 4 hours after adding labelled glucose to culture. (E-G) Analysis of the relative levels of glucose, G6P, 6PGL, 6PG, and S7P, and E4P levels in subcutaneous tumors formed by *G6PD* mutant and control A375 (*left*), M214 (*center*), and M481 (*right*) melanoma cells (n=5-8 tumors per clone). Each dot represents a different tumor from a different mouse. Data represent mean ±s.d. Statistical significance was assessed using linear mixed-effects analysis (B), linear mixed-effects analysis on log2-transformed data (F-H), Student’s t-tests (D), or Mann-Whitney tests (C). Statistical tests were two-sided. Multiple comparisons were adjusted by the FDR method. *P<0.05, **P<0.01, ***P<0.001, ****P<0.0001, ns= not significant.

To assess the effect of the hypomorphic *G6PD* mutations on pentose phosphate pathway function we performed isotope tracing by supplementing A375 cells with [1,2-^13^C] glucose in culture, then comparing the fractional enrichment in pentose phosphate pathway metabolites in *G6PD* mutant as compared to control clones. We observed little or no difference between *G6PD* mutant and control cells in the fractional enrichments (m+2) of glucose, glucose-6-phosphate, or in the non-oxidative pentose phosphate pathway intermediates sedoheptulose 7-phosphate (S7P) or erythrose 4-phosphate (E4P); however, the fractional enrichments in the oxidative pentose phosphate pathway intermediates 6-phosphogluconolactone (6PGL) and 6-phosphogluconate (6PG) were lower in *G6PD* mutant as compared to control clones (Fig. 2C). We also compared the relative flux of labelled glucose through glycolysis versus the oxidative pentose phosphate pathway by comparing the ratio of M+1 lactate (derived from the oxidative pentose phosphate pathway) to M+2 lactate (derived from glycolysis) (34). This ratio was significantly reduced in *G6PD* mutant as compared to control cells (Fig. 2D). Isotope tracing thus suggested that reduced G6PD function impaired the function of the oxidative branch of the pentose phosphate pathway, as expected.

We performed metabolomic analysis on primary subcutaneous tumors formed by *G6PD* mutant as compared to control melanoma cells. The levels of glucose and glucose-6-phosphate (Fig. 2E) and the non-oxidative pentose phosphate pathway intermediates S7P and E4P (Fig. 2G) generally did not significantly differ between *G6PD* mutant and control tumors. Consistent with the isotope tracing data, metabolites within the oxidative pentose phosphate pathway, 6PGL and 6PG, were almost always significantly depleted within *G6PD* mutant as compared to control tumors (Fig. 2F). Thus, both isotope tracing and metabolomics data suggest that *G6PD* mutant melanomas have diminished flux through the oxidative pentose phosphate pathway.

### Metastasizing melanoma cells depend on G6PD

To test whether G6PD deficiency affected subcutaneous tumor growth or metastasis, we subcutaneously transplanted 100 *G6PD* mutant or control melanoma cells into NSG mice. Nearly all of the injections of control cells and most of the injections of *G6PD* mutant cells formed tumors, though some clones of *G6PD* mutant A375 and M214 cells formed fewer tumors (Fig. 3A). The *G6PD* mutant and control A375 tumors grew at similar rates though the *G6PD* mutant M214 and M481 tumors grew more slowly than control tumors (Fig. 3B). Once tumors reached 2.5 cm, we assessed spontaneous metastasis by measuring the frequency of circulating melanoma cells in the blood and the metastatic disease burden using bioluminescence imaging of organs. The frequency of circulating melanoma cells in the blood was generally lower in mice with *G6PD* mutant as compared to control melanomas (Fig. 3C and Fig. S2B for flow cytometry gating strategy). Metastatic disease burden (Fig. 3D) and the percentage of mice that spontaneously formed macrometastases (Fig. 3E) was always significantly lower in mice with *G6PD* mutant as compared to control melanomas. In the case of A375, it is particularly striking that subcutaneous tumors formed by *G6PD* mutant cells grew at a similar rate as tumors formed by control cells and yet the frequency of circulating melanoma cells in the blood, overall metastatic disease burden, and the percentage of mice that formed macrometastases were all significantly lower in mice with *G6PD* mutant as compared to control cells. The data suggest that melanoma cells become more dependent upon G6PD during metastasis, consistent with the increase in oxidative stress during metastasis (15, 20, 21).

**Figure 3:**
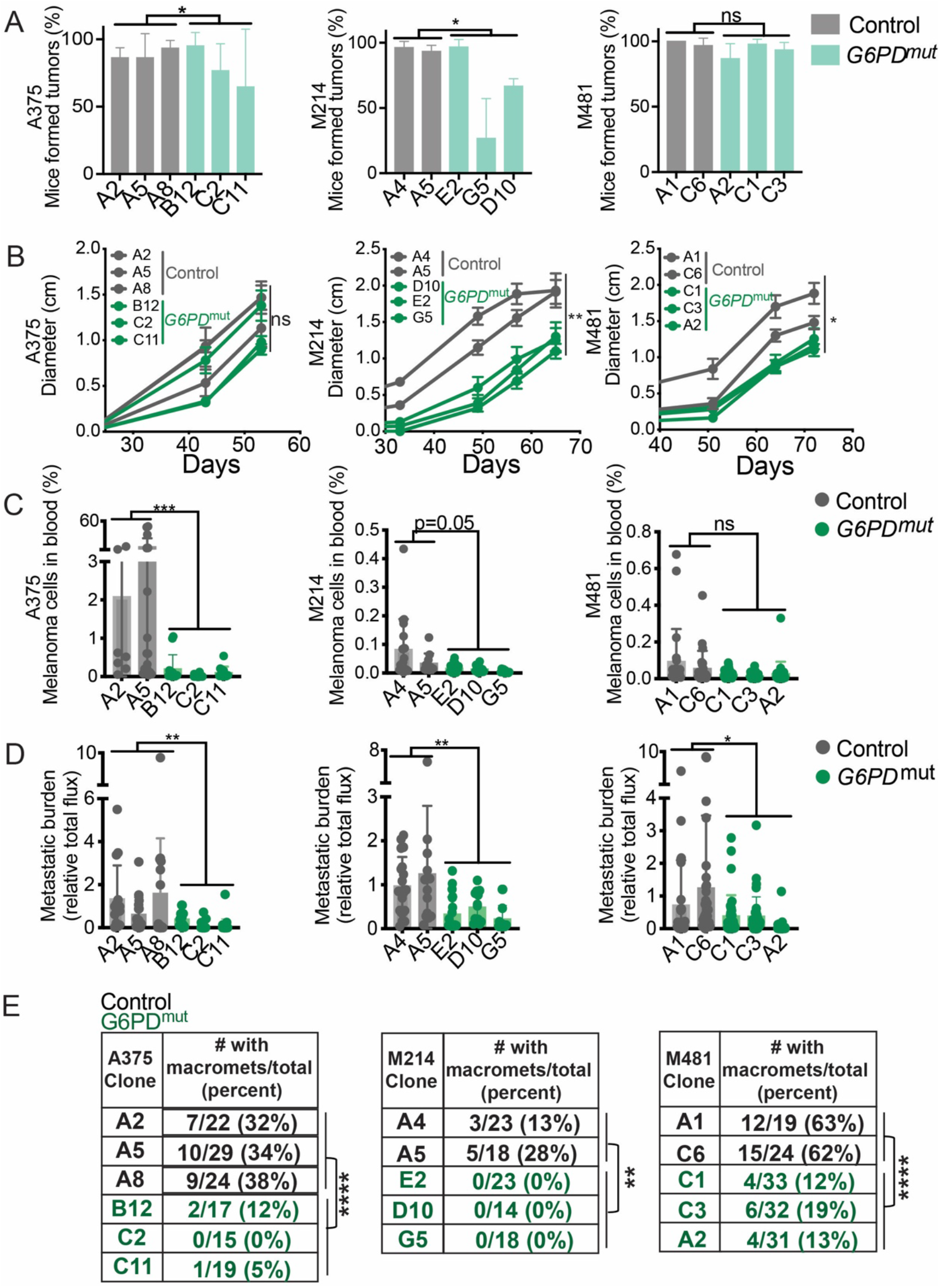
*G6PD* deficiency has limited effects on the growth of subcutaneous tumors but impairs metastasis. (A-E) NSG mice were injected subcutaneously (100 cells/injection) with *G6PD* mutant or control cells from A375, M214, and M481 melanomas and the tumors were allowed to spontaneously metastasize. (A) The percentage of mice that formed subcutaneous tumors. (B) The growth of subcutaneous tumors. (C) The frequency of circulating melanoma cells in the blood. (D) Metastatic disease burden by bioluminescence imaging. (E) The percentage of mice that formed macrometastases. Three independent experiments were performed per melanoma but panel B reflects results from a single representative experiment per melanoma due to the difficulty of combining tumor growth rate data from independent experiments. Data represent mean ±s.d. Statistical significance was assessed using generalized linear mixed-effects analysis (A, E), linear mixed-effects analysis (B), or linear mixed-effects analysis on log2-transformed data (C,D). Statistical tests were two-sided. Multiple comparisons were adjusted by the FDR method. *P<0.05, **P<0.01, ****P<0.0001, ns= not significant.

Levels of reactive oxygen species (ROS) tended to be significantly higher in subcutaneous tumors formed by *G6PD* mutant as compared to control melanoma cells based on both CellRox Green (Fig. 4A) and CellRox Red (Fig. 4B) staining. *G6PD* mutant melanomas consistently had lower NADPH/NADP+ ratios as compared to control cells (Fig. 4C), primarily due to a decrease in NADPH levels in the *G6PD* mutant melanomas. NADH/NAD+ ratios did not significantly differ except in A375 cells (Fig. 4D). The ratios of glutathione (GSH) to oxidized glutathione (GSSG) were significantly lower in *G6PD* mutant as compared to control melanomas (Fig. 4E). The data thus suggest that melanoma cells experienced higher levels of oxidative stress when G6PD function was decreased.

**Figure 4:**
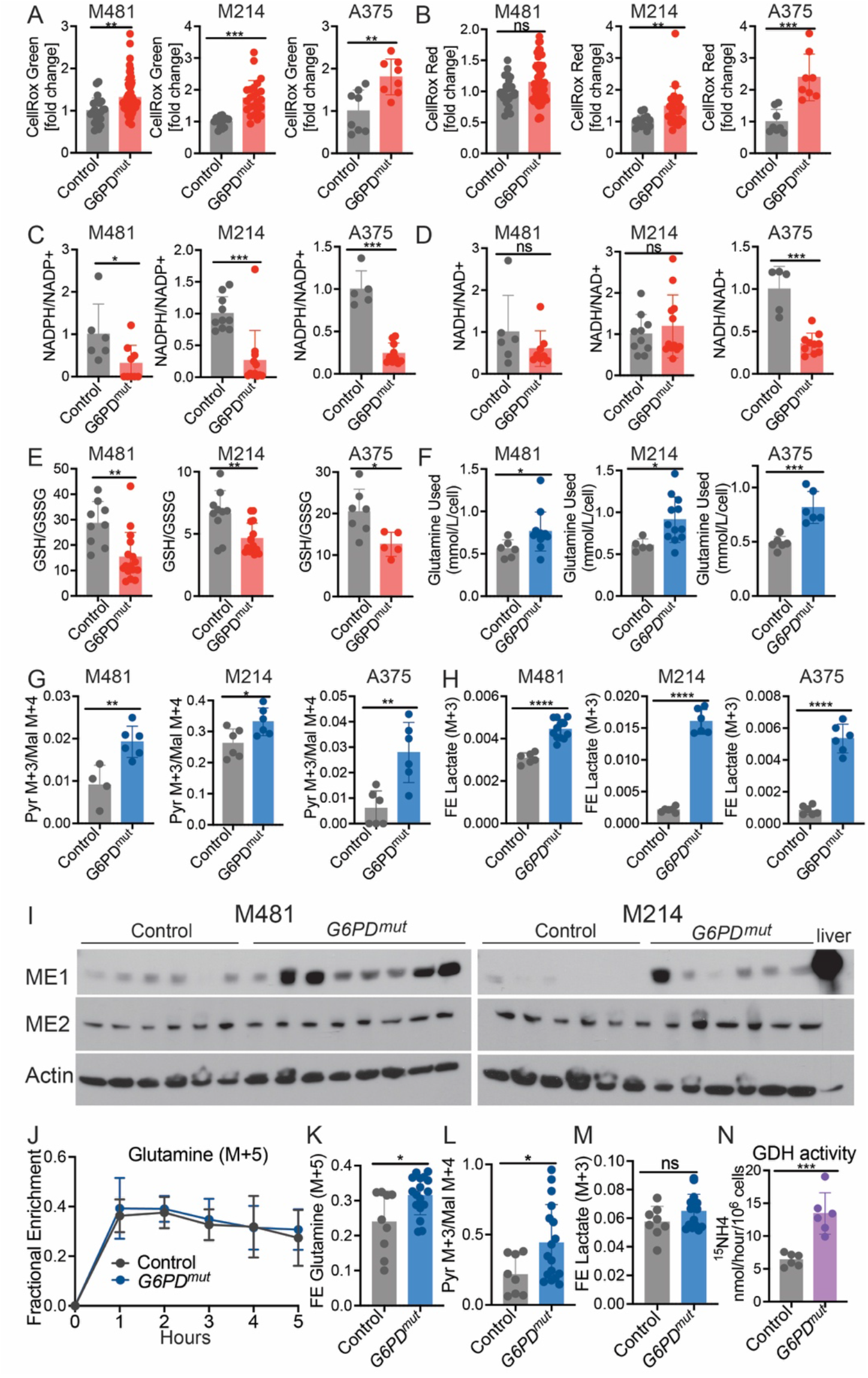
Reduced *G6PD* function induces oxidative stress and increases glutamine utilization. (A-B) ROS levels based on median fluorescence intensity of CellRox Green (*A*) or CellRox Red (*B*) in subcutaneous tumors formed by *G6PD* mutant or control melanoma cells (n=2-3 clones per genotype per melanoma with 5 mice per clone in 2-3 independent experiments per melanoma). Each dot represents data from one mouse. (C-E) The relative ratios of NADPH to NADP+ (*C*) and NADH to NAD+ (*D*) and the absolute molar ratios of GSH to GSSG quantified with an internal standard (*E*) in *G6PD* mutant or control M481, M214 and A375 melanoma cells (three independent experiments were performed per melanoma and one representative experiment per melanoma is shown; n= 5-15 mice per group). (F) The amount of glutamine consumed from the culture medium is shown as mmol/liter/cell for *G6PD* mutant and control melanomas after 24 (M481 and M214) or 8 hours (A375) in culture (n= 2 clones per genotype per melanoma, with 3-6 replicate cultures per clone in 2 independent experiments per melanoma). (G-H) *G6PD* mutant or control melanoma cells were cultured with [U-^13^C] glutamine. The ratio of pyruvate m+3 to malate m+4 reflects malic enzyme activity (*G*) and fractional enrichment in lactate (m+3) reflects the relative rate of reduction of pyruvate by lactate dehydrogenase (*H*). (I) Western blot analysis of Malic Enzyme 1 (ME1) and Malic Enzyme 2 (ME2) in lysates of subcutaneous tumors formed by *G6PD* mutant and control M481 (*left*) or M214 (*right*) melanomas. Actin is a loading control. Each lane is a different tumor from a different mouse. Normal mouse liver is shown as a positive control. (J-M) NSG mice xenografted with *G6PD* mutant or control M481 melanoma cells were infused with [U-^13^C] glutamine. The fractional enrichment of glutamine m+5 in the blood (*J*) and in subcutaneous tumors after 5 hours of infusion (*K*). The ratio of pyruvate m+3 to malate m+4 (*L*) and fractional enrichment in lactate (m+3) (*M*) (n= 2 clones per genotype per melanoma, with 4-8 mice per clone in 2 independent experiments). Each dot represents a different tumor in a different mouse. (N) *G6PD* mutant and control melanoma cells were cultured in medium supplemented with L-[α-^15^N]glutamine and GDH activity was measured based on the rate at which ^15^N was transferred from glutamine to ^15^NH_4_ (the data represent 2 clones per genotype from three independent experiments; each dot represents cells from a different culture; n=3 wells/clone). Data represent mean ±s.d. Statistical significance was assessed using Mann-Whitney tests (A-C, J, and M), Student’s t-tests on log2-transformed data (D-F), Student’s t-tests (G-H, K-M), Welch’s t-test (H), or Student’s t-tests on log2-transformed data (N). Statistical tests were two-sided. Multiple comparisons were adjusted by the FDR method: *P<0.05, **P<0.01, ***P<0.001, ****P<0.0001, ns=not significant.

### G6PD mutant melanomas increase glutaminolysis

We wondered whether *G6PD* mutant melanomas might compensate for the loss of G6PD function by increasing NADPH production through other mechanisms. *G6PD* deficient colon cancer cells upregulate malic enzyme flux relative to control cells based on isotope tracing from labelled glutamine (1). We first compared the consumption of glutamine by *G6PD* mutant and control melanoma cells in culture. We found that the *G6PD* mutant cells consistently consumed more glutamine from the culture medium than control cells (Fig. 4F).

To test if *G6PD* mutant melanomas compensate by increasing glutaminolysis (the conversion of glutamine to pyruvate through the TCA cycle, with the production of NADPH by malic enzyme; Fig. S1), we performed isotope tracing with [U-^13^C] glutamine in culture. [U-^13^C] glutamine gives rise to 4-carbon TCA cycle intermediates, such as malate (m+4). Because malic enzyme decarboxylates malate to produce pyruvate, malic enzyme activity is reflected in the ratio of pyruvate m+3 to malate m+4. This ratio was consistently and significantly higher in *G6PD* mutant as compared to control melanoma cells (Fig. 4G). The fractional enrichment of (m+3) lactate was also significantly higher in *G6PD* mutant as compared to control melanoma cells (Fig. 4H). Malic enzyme 2 (ME2), which localizes to mitochondria, was expressed by all melanomas at similar levels in *G6PD* mutant as compared to control cells (Fig. 4I). Malic enzyme 1 (ME1), which localizes to the cytoplasm, was expressed by most melanomas but it tended to be at higher levels in *G6PD* mutant as compared to control melanomas (Fig. 4I). These data suggest that *G6PD* mutant melanomas compensate for the loss of G6PD function by increasing glutaminolysis and malic enzyme activity, partly by increasing ME1 levels.

To assess this in vivo, we infused [U-^13^C] glutamine into mice xenografted with *G6PD* mutant or control M481 melanomas (35). Infusion of [U-^13^C] glutamine enriched the circulating glutamine pool to a similar extent in mice transplanted with *G6PD* mutant and control melanomas (Fig. 4J). Consistent with the results in culture, the fractional enrichment of (m+5) glutamine was significantly higher in *G6PD* mutant as compared to control melanomas (Fig. 4K). Malic enzyme activity, based on the ratio of glutamine derived pyruvate (m+3) to glutamine derived (m+4) malate, was significantly higher in *G6PD* mutant as compared to control melanoma cells (Fig. 4L). This ratio was higher in vivo than in the same melanoma cells cultured with glutamine (m+5) in vitro (compare Fig. 4L and Fig. 4G, *left panel*), regardless of *G6PD* status. This difference likely results in part from glutaminolysis and gluconeogenesis occurring outside the tumor. These pathways can produce circulating glucose m+3 and lactate m+3 from [U-^13^C] glutamine and thereby indirectly contribute to pyruvate m+3 in the tumor (36). Nevertheless, the elevated pyruvate m+3 to malate m+4 ratio in *G6PD* mutant melanomas, as compared to control melanomas, is consistent with a relative increase in malic enzyme’s contribution to the pyruvate pool in the *G6PD* mutant tumors, as observed in culture. The fractional enrichment of (m+3) lactate also tended to be higher in *G6PD* mutant as compared to control melanoma cells, though the difference was not statistically significant (Fig. 4M). These data suggest that *G6PD* mutant melanomas increase malic enzyme activity in vivo.

Cells convert glutamine into the TCA cycle intermediate α-ketoglutarate (αKG) via glutaminase (GLS) and glutamate dehydrogenase (GDH) (Fig. S1). GDH converts glutamate to αKG while generating ammonia (NH_3_) from the a nitrogen of glutamine. To test if *G6PD* mutant melanomas exhibit higher GDH activity we cultured cells with ^15^N-glutamine (labeled in the aposition) and measured the production of ^15^NH_4_^+^ by GC/MS (37). GDH activity was significantly increased in *G6PD* mutant as compared to control melanomas (Fig. 4N). This suggests *G6PD* mutant melanomas increase both GDH and malic enzyme activity to increase the generation of reducing equivalents in the absence of oxidative pentose phosphate pathway function.

### *G6PD* deficiency sensitizes melanoma cells to glutaminase inhibition

Glutamine dependency has been targeted in several cancers, primarily by inhibiting glutaminase (38). To test whether *G6PD* mutant melanoma cells were sensitive to glutaminase inhibition we cultured *G6PD* mutant or control melanoma cells in vehicle (DMSO) or CB-839, a glutaminase inhibitor (39). 1nM CB-839 did not significantly affect the growth of A375 control melanoma cells and 10nM CB-839 only modestly reduced their growth (Fig. 5A). In contrast, 1nM and 10nM CB-839 completely blocked the growth of *G6PD^mut^* A375 cells (Fig. 5B). No concentration of CB-839 reduced the growth of M214 control cells (Fig. 5C); however, all concentrations of CB-839 (1 to 1000 nM) significantly reduced the growth of *G6PD^mut^* M214 cells (Fig. 5D). *G6PD^mut^* melanoma cells were therefore far more sensitive to glutaminase inhibition than control melanoma cells.

**Figure 5:**
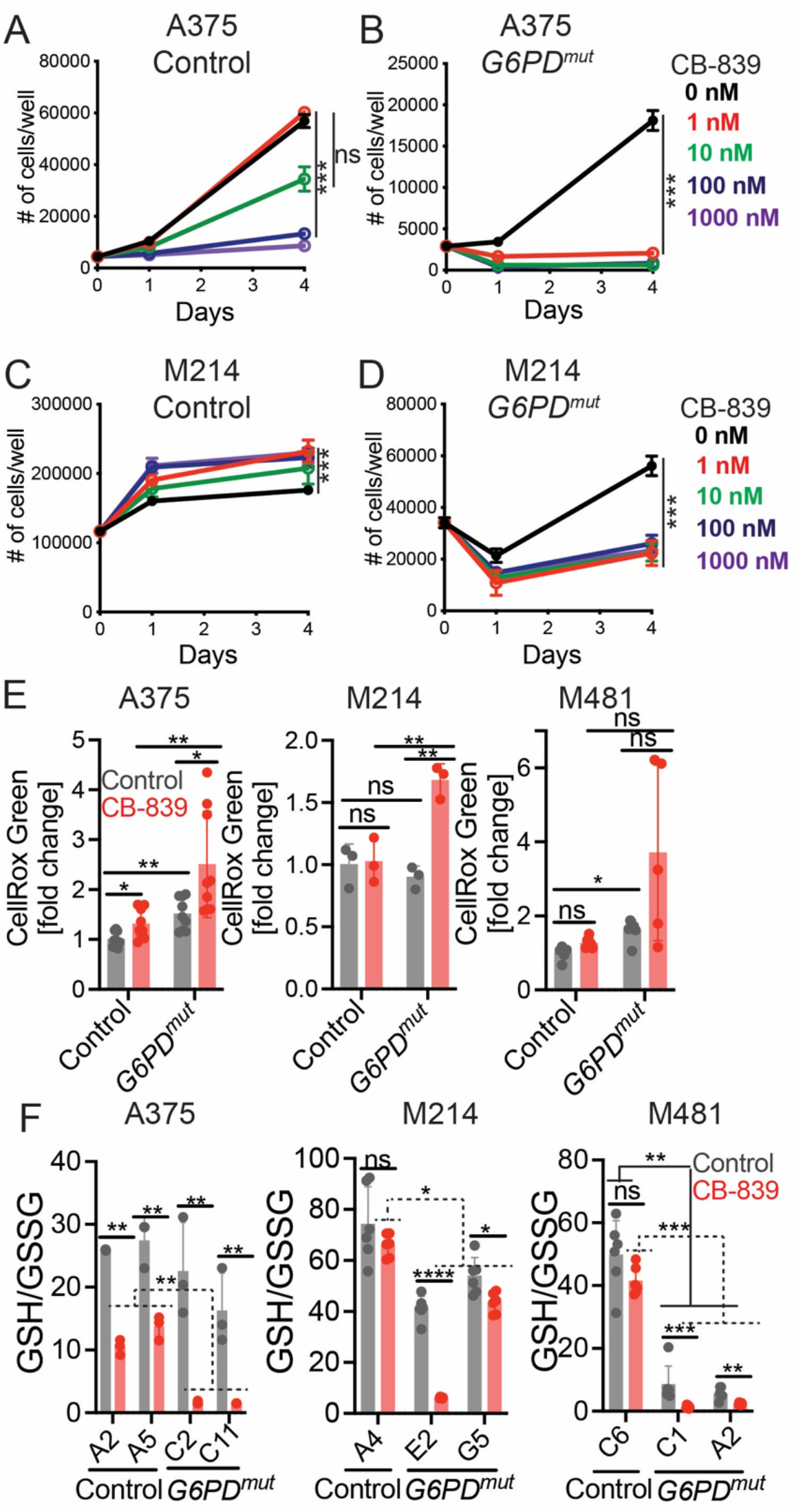
*G6PD* deficiency sensitizes melanoma cells to glutaminase inhibition. (A-D) *G6PD* mutant and control A375 (*A-B*) or M214 (*C-D*) melanoma cells were cultured with the glutaminase inhibitor, CB-839 (1 nM, 10 nM, 100 nM, or 1000 nM), or vehicle (DMSO) control (0 nM for CB-839) and the number of cells in each well was counted over time. (E-F) A375, M214, or M481 melanoma cells were cultured in the presence of CB-839 (100 nM) or vehicle for 24 hours and ROS levels were analyzed based on flow cytometric analysis of CellRox Green staining (E) and the absolute molar ratios of GSH to GSSG were quantified by mass spectrometry with an internal standard (F). Each panel reflects data from a single experiment that is representative of two to three independent experiments, with 1-2 clones per genotype and 3-6 replicate cultures per clone per experiment. All data represent mean ±s.d. Statistical significance was assessed using nparLD (A-D), Welch’s one-way ANOVA with log2-transformed data followed by Dunnett’s T3 test for multiple comparisons adjustment (E), or Welch’s t-tests with log2-transformed data (F). Statistical tests were two-sided. Multiple comparisons were adjusted by the FDR method except for panel (E): *P<0.05, **P<0.01, ***P<0.001,****P<0.0001, ns=not significant.

To test if glutaminase inhibition increased oxidative stress in *G6PD^mut^* melanoma cells, we measured ROS levels (CellRox Green) and GSH to GSSG ratios in *G6PD* mutant and control melanoma cells treated in culture with 100nM CB-839 or vehicle control. CB-839 modestly increased ROS levels in A375 control cells but not in M214 or M481 control cells (Fig. 5E). ROS levels were usually higher in *G6PD* mutant as compared to control melanoma cells (including A375 and M481, but not M214; Fig. 5E). ROS levels were highest in *G6PD* mutant A375, M214, and M481 cells treated with CB-839, generally higher than in untreated *G6PD* mutant cells or in CB-839 treated control cells (Fig. 5E). Glutaminase inhibition thus increased ROS levels to a greater extent in *G6PD* mutant as compared to control melanoma cells.

CB-839 treatment significantly reduced GSH/GSSG ratios in A375 control cells but not in M214 or M481 control cells (Fig. 5F). *G6PD* deficiency also reduced GSH/GSSG ratios in A375 control cells but not in M214 or M481 control cells (Fig. 5F). GSH/GSSG ratios were substantially lower in *G6PD* mutant A375, M214, and M481 cells treated with CB-839 as compared to either untreated *G6PD* mutant cells or CB-839 treated control cells (Fig. 5F). Glutaminase inhibition typically reduced GSH/GSSG ratios to a much greater extent in *G6PD* mutant as compared to control melanoma cells. Taken together, the data suggest that when the oxidative pentose phosphate pathway is impaired by reduced G6PD function, melanoma cells become more dependent upon glutaminolysis for oxidative stress management.

## Discussion

Our data suggest that melanoma cells become more dependent upon the oxidative pentose phosphate pathway during metastasis. Melanoma cells experience a spike in ROS during metastasis, leading to oxidative stress and the death of most metastasizing cells by ferroptosis (15, 16, 21, 40). This is observed in both patient-derived xenografts growing in immunocompromised mice as well as mouse melanomas growing in syngeneic, immunocompetent mice. Oxidative stress limits the metastasis of other cancers as well (41–50), though oxidative stress may promote metastasis under certain circumstances (51, 52). Large clinical trials found that antioxidant supplementation in humans increases cancer incidence and mortality (53–57). Pro-oxidant therapies have the potential to inhibit cancer progression by exacerbating the oxidative stress experienced by metastasizing cancer cells and nascent metastatic nodules (17).

To develop effective pro-oxidant therapies, it will be necessary to identify the diverse mechanisms cancer cells use to manage oxidative stress during metastasis. Our results show that melanoma cells have redundant and layered protection against oxidative stress: reduced G6PD function led to compensatory upregulation of glutaminolysis and malic enzyme activity. This is consistent with results in other cancers, which exhibited compensatory increases in isocitrate dehydrogenase activity, malic enzyme activity, and folate metabolism in response to reduced G6PD function (1, 26). Consequently, reduced G6PD function decreases, but does not completely eliminate, metastasis.

## Materials and methods

### Melanoma specimens and enzymatic dissociation

Melanoma specimens were obtained with informed consent from patients according to protocols approved by the Institutional Review Board of the University of Michigan Medical School (IRBMED approvals HUM00050754 and HUM0005008524) and the University of Texas Southwestern Medical Center (IRB approval 102010-051). Materials used in the manuscript are available, either commercially or from the authors, though there are restrictions imposed by Institutional Review Board requirements and institutional policy on the sharing of materials from patients. Single-cell suspensions were obtained by dissociating tumors in Kontes tubes with disposable pestles (VWR) followed by enzymatic dissociation in 200 U/ml collagenase IV (Worthington), DNase (50 U/ml) and 5 mM CaCl_2_ at 37 °C for 20 min. We typically used 1 to 5 ml of dissociation medium, depending on the size of the tumors. After dissociation, cell suspensions were filtered through a 40 μm cell strainer to remove clumps.

### Mouse studies and xenograft assays

All mouse experiments complied with all relevant ethical regulations and were performed according to protocols approved by the Institutional Animal Care and Use Committee at the University of Texas Southwestern Medical Center (protocol 2016-101360). Melanoma cell suspensions were prepared for injection in staining medium (L15 medium containing bovine serum albumin (1 mg/ml), 1% penicillin/streptomycin and 10 mM HEPES (pH 7.4) with 25% high-protein Matrigel (product 354248; BD Biosciences)). Subcutaneous injections were performed in the right flank of NOD.CB17-*Prkdc^scid^ Il2rg^tm1Wjl^*/SzJ (NSG) mice in a final volume of 50 μl. Four-to-eight-week-old male and female NSG mice were transplanted with 100 melanoma cells, subcutaneously unless otherwise specified. Both male and female mice were used. Subcutaneous tumor diameters were measured weekly with calipers until any tumor in the mouse cohort reached 2.5 cm in its largest diameter, in agreement with the approved animal protocol. At that point, all mice in the cohort were euthanized and spontaneous metastasis was evaluated by gross inspection of visceral organs for macrometastases and bioluminescence imaging of visceral organs to quantify metastatic disease burden (see details below).

### Bioluminescence imaging

All melanomas were tagged with stable luciferase expression, enabling the quantitation of metastatic disease burden by bioluminescence imaging. Five minutes before performing luminescence imaging, mice were injected intraperitoneally with 100 μl of PBS containing D-luciferin monopotassium salt (40 mg/ml) (Goldbio) and mice were anaesthetized with isoflurane 2 min before imaging. All mice were imaged using an IVIS Lumina S5 (Perkin Elmer) with Living Image software (Perkin Elmer). After completion of whole-body imaging, mice were euthanized and individual organs (including the heart, lung, liver, pancreas, spleen, and kidney) were surgically removed and imaged. The exposure time ranged from 15 to 30 seconds, depending on the maximum signal intensity, to avoid saturation of the luminescence signal. To measure the background luminescence, a negative control mouse not transplanted with melanoma cells was imaged. The bioluminescence signal (total photon flux) was quantified with ‘region of interest’ measurement tools in Living Image software. Metastatic disease burden was calculated as observed total photon flux from all organs in xenografted mice minus background total photon flux in negative control mice. Negative values were set to 1 for purposes of presentation and statistical analysis.

### Cell labelling and flow cytometry

Melanoma cells were identified and sorted by flow cytometry as previously described (15, 20). All antibody staining was performed for 20 min on ice, followed by washing with HBSS and centrifugation at 200xg for 5 min. Cells were stained with directly conjugated antibodies against mouse CD45 (APC, eBiosciences), mouse CD31 (APC, Biolegend), mouse Ter119 (APC, eBiosciences) and human HLA-ABC (G46-2.6-FITC, BD Biosciences). Human melanoma cells were isolated as cells that were positive for HLA and negative for mouse endothelial and hematopoietic markers. Cells were washed with staining medium and re-suspended in 4’,6-diamidino-2-phenylindole (DAPI; 1 μg/ml; Sigma) to eliminate dead cells from sorts and analyses. Cells were examined on an LSRFortessa cell analyzer (Becton Dickinson) or sorted on a FACS Fusion Cell Sorter (Becton Dickinson). For analysis of circulating melanoma cells, blood was collected from mice by cardiac puncture with a syringe pretreated with citrate-dextrose solution (Sigma) when subcutaneous tumors reached 2.5 cm in diameter. Red blood cells were sedimented using Ficoll, according to the manufacturer’s instructions (Ficoll Paque Plus, GE Healthcare). Remaining cells were washed with HBSS (Invitrogen) before antibody staining and flow cytometry.

### CRISPR editing of *G6PD* in melanoma cells

Single-guide RNAs (sgRNAs) targeting exon 6 of human *G6PD* were designed using publicly available tools (http://crispr.mit.edu): *G6PD* sgRNA #1, 5’–CTGGTCCTCACGGAACAGGG–3’; *G6PD* sgRNA #2, 5’–AGGCTGGAACCGCATCATCG–3’. The sgRNAs were cloned into the U6-driven Cas9 expression vector (pX458-pSpCas9(BB)-2AGFP; 48318, Addgene) (58). sgRNA insertion was confirmed by Sanger sequencing.

Approximately 100,000 to 500,000 melanoma cells were plated in tissue-culture-treated 6-well plates in DMEM plus 10% FBS and 1% penicillin/streptomycin. 1 μg of each of the two sgRNA constructs was co-transfected into the melanoma cells using polyjet (SignaGen) according to the manufacturer’s instructions. After 36–48 hours, GFP+ cells were flow cytometrically isolated and allowed to recover for 24–48 hours in DMEM plus 10% FBS and 1% penicillin/streptomycin. Single cells were then plated in tissue-culture-treated 96-well plates in Prime-XV tumorsphere medium (Irvine Scientific) supplemented with 2 U/ml Heparin (Sigma), 0.1 μg/ml Hydrocortisone (Sigma), 2% B27 (Thermo Fisher), 1 μM StemRegenin 1 (StemCell Technologies, Inc.), 10% charcoal stripped FBS (Thermo Fisher), 10 μg/ml Bovine Pituitary Extract (BPE; Lonza), 10 ng/ml recombinant human IL-8 (CXCL8; Peprotech), 20 ng/ml recombinant human GRO-α/MGSA (CXCL1; Peprotech), and 25 ng/ml recombinant human HGF (Peprotech). Genomic DNA was isolated from individual clones with QuickExtract (Epicenter) and clones were screened and sequenced for *G6PD* deletions.

### G6PD activity assay

G6PD enzyme activity was determined using the Glucose-6-Phosphate Dehydrogenase (G6PD) Activity Assay Kit (12581S, Cell Signaling Technology) according to the manufacturer’s instructions. Lysates were collected from subcutaneous melanomas or melanoma cells growing in culture and added to assay buffer. Glucose-6-phosphate (G6P), in the presence of NADP, is oxidized by G6PD to generate 6-phosphogluconolactone and NADPH. The NADPH is then amplified by the diaphorase-cycling system to produce fluorescent resorufin molecules. The relative fluorescent units (RFU) was determined using a plate reader with excitation at 540 nm and emission at 590 nm. RFU is proportional to G6PD activity in this assay.

### Metabolomic analysis

HILIC chromatographic separation of metabolites was achieved using a Millipore ZIC-pHILIC column (5 μm, 2.1 × 150 mm) with a binary solvent system of 10 mM ammonium acetate in water, pH 9.8 (solvent A) and acetonitrile (solvent B) with a constant flow rate of 0.25 ml/minute. For gradient separation, the column was equilibrated with 90% solvent B. After injection, the gradient proceeded as follows: 0–15 min linear ramp from 90% B to 30% B; 15–18 min isocratic flow of 30% B; 18–19 min linear ramp from 30% B to 90% B; 19–27 column regeneration with isocratic flow of 90% B. Metabolites were measured with a Thermo Scientific QExactive HF-X hybrid quadrupole orbitrap high-resolution mass spectrometer (HRMS) coupled to a Vanquish UHPLC. HRMS data were acquired with two separate acquisition methods. Individual samples were acquired with an HRMS full scan (precursor ion only) method switching between positive and negative polarities. For data-dependent, high-resolution tandem mass spectrometry (ddHRMS/MS) methods, precursor ion scans were acquired at a resolving power of 60,000 full width at half-maximum (FWHM) with a mass range of 80–1,200 Da. The AGC target value was set to 1 × 10^6^ with a maximum injection time of 100 ms. Pooled samples were generated from an equal mixture of all individual samples and analyzed using individual positive- and negative-polarity spectrometry ddHRMS/MS acquisition methods for high-confidence metabolite identification. Product ion spectra were acquired at a resolving power of 15,000 FWHM without a fixed mass range. The AGC target value was set to 2 × 10^5^ with a maximum injection time of 150 ms. Data-dependent parameters were set to acquire the top 10 ions with a dynamic exclusion of 30 s and a mass tolerance of 5 ppm. Isotope exclusion was turned on and a normalized collision energy of 30 was applied. Settings remained the same in both polarities.

Metabolite identities were confirmed in three ways: (1) precursor ion m/z was matched within 5 ppm of theoretical mass predicted by the chemical formula; (2) fragment ion spectra were matched within a 5 ppm tolerance to known metabolite fragments; and (3) the retention time of metabolites was within 5% of the retention time of a purified standard run with the same chromatographic method. Metabolites were relatively quantitated by integrating the chromatographic peak area of the precursor ion searched within a 5 ppm tolerance. To analyze low-abundance metabolites extracted from tumors (such as NADPH, NADP+, NAD+, NADH, 6-phosphogluconate, and 6-phosphogluconolactone), we created an additional targeted selected ion monitoring (tSIM) scan. We used a resolving power of 60,000 FWHM and an AGC target of 1 x 10^5^ with a maximum injection time of 100 ms for these scan events. An inclusion list for each m/z of interest was created with an isolation window of 5 daltons and an isolation offset of 1 dalton. NADPH/NADP+ ratios were determined by integrating the extracted ion chromatograms for NADPH in the negative tSIM scan (*m*/*z* = 744.0838) and NADP^+^ in the positive tSIM scan (*m*/*z* = 744.0827). Fragmentation spectra from pooled samples were used for structural confirmation of NADPH and NADP^+^.

For analysis of the GSH to GSSG ratio by liquid chromatography– tandem mass spectrometry (LC–MS/MS), subcutaneous tumor fragments weighing 5–15 mg were homogenized using Kontes tubes with disposable pestles (VWR) in ice-cold 80:20 methanol:water (v/v), with 0.1% formic acid to prevent spontaneous oxidation (59), followed by three freeze-thaw cycles in liquid nitrogen. The supernatant was collected after a 10-min centrifugation at 13,000xg at 4°C then lyophilized. Lyophilized samples were reconstituted in 100 μl of 0.1% formic acid in water, vortexed and analyzed by LC–MS/MS. GSH/GSSG analysis was performed using a SCIEX 6500+ Q-Trap mass spectrometer coupled to a Shimadzu LC-20A UHPLC system. Chromatographic separation was carried out with a Waters HSS T3 column and a binary solvent gradient of water with 0.1% formic acid (solvent A) and acetonitrile with 0.1% formic acid (solvent B). The following gradient was used for separation: 0–3 min, isocratic flow of 0% B; 3–8 min, 0–100% B; 8–13 min, isocratic flow of 100% B; 13–13.1 min, 100–0% B; 13.1–18 min, isocratic flow of 0% B. The flow rate was held constant at 0.2 ml/minute. The mass spectrometer was operated in MRM mode, monitoring the following transitions for GSH, GSSH and their respective internal standards in positive mode: GSH 308/162; GSSG 613/355; GSH internal standard (ISTD) 311/165; GSSG ISTD 619/165. Transitions and source parameters were optimized by infusion before analysis. GSH/GSSG ratios were calculated based on the molar values of GSH and GSSG, determined using a standard curve and internal standards.

### Isotope tracing

We extracted and acquired mass spectra of metabolites from isotopically labeled specimens using the same methods described above. Analysis of fractional enrichment was achieved by calculating the theoretical mass of all ^13^C isotopologues of a metabolite and integrating the resulting peak in the extracted ion chromatogram. We used the following criteria to ensure the correct peaks were integrated for analysis: (1) the precursor ion m/z of the M+0 peak was matched within 5 ppm of the theoretical mass predicted by its chemical formula; (2) the retention time of the M+0 peak was within 5% of the retention time of a purified chemical standard run with the same chromatographic method; (3) all isotopes of a potentially labeled metabolite were within 5 ppm of their predicted m/z by chemical formula; and (4) all isotopologues eluted simultaneously with the M+0 peak. An unlabeled sample was run alongside all isotopically labeled samples to acquire product ion spectra for additional verification. After analyzing the raw data, peak areas were further analyzed using a published algorithm to correct for naturally-abundant isotopes and calculate fractional enrichment (60).

To trace isotopically labelled glucose or glutamine in culture, melanoma cells were grown adherently in 6-well plates with DMEM lacking glucose, glutamine, and phenol red (A14430, Gibco), supplemented with 10% dialyzed fetal calf serum, 12.5 mM glucose, and penicillin/streptomycin. At time 0, the cells were washed with PBS and fed 1 ml of culture medium supplemented with 2 mM [U-^13^C]glutamine (CLM-1822, Cambridge Isotope Laboratories) or 12.5 mM [1,2-^13^C]glucose (CLM-504-1, Cambridge Isotope Laboratories). Lysates were harvested in 80% methanol at the time points indicated in the figures and analyzed by LC–MS/MS.

In vivo isotope tracing experiments were performed when subcutaneous tumors reached 1.5 to 2 cm in diameter. Before infusions, mice were fasted for 16 hours, then a 27-gauge catheter was placed in the lateral tail vein under anesthesia. We intravenously infused [U-^13^C]glutamine (CLM-1822, Cambridge Isotope Laboratories) as a bolus of 0.1725 mg/g body mass over 1 min in 150 μl of saline, followed by continuous infusion of 0.00288 mg/g body mass/minute for 5 hours in a volume of 150 μl/hour (36). For infusions of [1,2-^13^C]glucose (CLM-504, Cambridge Isotope Laboratories), we intravenously infused a bolus of 0.4125 mg/g body mass over 1 minute in 125 μl of saline, followed by continuous infusion of 0.008 mg/g body mass/min for 3 hours in a volume of 150 μl/hour (35). At the end of the infusion, mice were euthanized and tumors were collected and immediately frozen in liquid nitrogen. To assess the fractional enrichments in plasma, 20 μl of blood was obtained after 30, 60, 120 and 180 min of infusion (glucose) or hourly for 5 hours after infusion (glutamine).

### Glutaminase inhibitor treatment

To assess the growth of melanoma cells in the presence of the glutaminase inhibitor, CB-839 (Telaglenastat, S7655, Selleck Chemicals), melanoma cells were grown adherently in 24-well or 96-well plates. To test the effects of glutaminase inhibition, melanoma cells were cultured in DMEM + 10% FBS and at day 0 treated with 100 nM CB-839 or DMSO control. Cells were trypsinized at the indicated time points and the number of cells was counted manually using a hemocytometer or using an LSRFortessa flow cytometer. Dead cells were identified as trypan blue or DAPI positive and were excluded from the counts.

### Glutamine uptake and glutamate dehydrogenase activity

Glutamine concentrations in the culture medium were measured using an automated chemical analyzer (NOVA Biomedical). To determine the rate of glutamine consumption by cells, we measured the concentration of glutamine in the culture medium after the cells had been growing for 8-24 hours and compared it to the concentration of glutamine in culture medium incubated for the same period of time in wells without cells.

Glutamate dehydrogenase (GDH) activity in cultured melanoma cells was quantitated by measuring the transfer of ^15^N from [α–^15^N]glutamine (NLM-1016-1, Cambridge Isotope Laboratories) to ^15^NH_4_^+^ as described (37) after culture for 8 hours in the presence of [α-^15^N]glutamine. Total NH_4_^+^ secreted into the medium was measured by spectrophotometry (Megazyme). The fractional enrichment of ^15^N in NH_4_^+^ in the culture medium was determined using a modification of a published method (61): the NH_4_^+^ in the culture medium was conjugated to a ketoacid (ketovaleric acid) in the presence of purified glutamate dehydrogenase (GDH) and NADH for 30 min at 37°C to form an amino acid, α-aminobutyrate. The α-aminobutyrate was then derivatized by addition of TMS groups and the samples were purified by applying to an ion exchange column (AG 50W-X4, Bio-Rad), washed with water, and eluted with 2 ml of 4 N ammonium hydroxide. Standards containing known ratios of unlabeled/^15^N-labeled NH_4_^+^ were prepared using the same method. All samples were evaporated and derivatized in 100 μl of Tri-Sil (Thermo Scientific), then analyzed by gas chromatography/mass spectrometry (GC/MS) with an Agilent 6890N GC coupled to an Agilent 5973 Mass Selective Detector. The oven temperature was ramped from 70°C to 150°C by 5°C/min, then by 10°C/min to 325°C. We measured the ratio of aminobutyrate *m*/*z* 130 (unenriched) to 131 (enriched), corresponding to the unlabeled/^15^N-labeled NH_4_^+^ ratio, and then multiplied by the total nMoles of NH_4_^+^ in the medium to determine the molar amount of NH_4_^+^ produced by the cells via GDH.

### Analysis of ROS levels by flow cytometry

Subcutaneous tumors were surgically excised as quickly as possible after euthanizing the mice and then melanoma cells were mechanically dissociated in 700 μl of staining medium. Single cell suspensions were obtained by passing the dissociated cells through a 40 μm cell strainer. Equal numbers of dissociated cells from each tumor were stained for cell surface markers and then stained for 30 min at 37°C with 5 mM CellROX Green or CellROX DeepRed (Life Technologies) in HBSS-free (Ca^2+^ and Mg^2+^-free) with DAPI to distinguish live from dead cells. The cells were then washed and analyzed by flow cytometry using either a FACS Fusion or a LSRFortessa (BD Biosciences) to assess ROS levels in live human melanoma cells (positive for human HLA and dsRed and negative for DAPI and mouse CD45/CD31/Ter119).

### Western blot analysis

Melanomas were excised and quickly snap-frozen in liquid nitrogen. Tumor lysates were prepared in Kontes tubes with disposable pestles using RIPA Buffer (Cell Signaling Technology) supplemented with phenylmethylsulphonyl fluoride (Sigma), and protease and phosphatase inhibitor cocktail (Roche). The bicinchoninic acid protein assay (Thermo) was used to quantify protein concentrations. Equal amounts of protein (5–20 μg) were loaded into each lane and separated on 4–20% polyacrylamide tris glycine SDS gels (BioRad), then transferred to polyvinylideneifluoride membranes (BioRad). The membranes were blocked for 1 h at room temperature with 5% milk or 5% BSA in TBS supplemented with 0.1% Tween-20 (TBST) and then incubated with primary antibodies overnight at 4°C. After washing, then incubating with horseradish peroxidase conjugated secondary antibody (Cell Signaling Technology), signals were developed using SuperSignal West (Thermo Fisher). Blots were sometimes stripped using Restore stripping buffer (Thermo Fisher) and re-stained with other primary antibodies. The following antibodies were used for western blots: anti-G6PD (D5D2, Cell Signaling Technologies), anti-ME1 (PA5-21550, Thermo Scientific), anti-ME2 (12399, Cell Signaling Technologies), anti-GAPDH (14C10, Cell Signaling Technologies), and anti--β-actin (D6A8, Cell Signaling Technologies).

### Statistical methods

Figures generally reflect data obtained in multiple independent experiments performed using different mice or cultures on different days. Mice were allocated to experiments randomly and samples processed in an arbitrary order, but formal randomization techniques were not used. Before analyzing the statistical significance of differences among treatments, we tested whether data were normally distributed and whether variance was similar among treatments. To test for normality, we performed the Shapiro–Wilk tests when 3 ≤ n < 20 or D’Agostino omnibus tests when n ≥ 20. To test whether variability significantly differed among treatments we performed F-tests (for experiments with two treatments) or Levene’s median tests (for experiments with more than two treatments). When the data significantly deviated from normality (P < 0.01) or variability significantly differed among treatments (P < 0.05), we log2-transformed the data and tested again for normality and variability. If the transformed data no longer significantly deviated from normality and equal variability, we performed parametric tests on the transformed data. If log2-transformation was not possible or the transformed data still significantly deviated from normality or equal variability, we performed non-parametric tests on the non-transformed data.

All of the statistical tests we used were two-sided. To assess the statistical significance of a difference between two treatments, we used Student’s t-tests (when a parametric test was appropriate), Welch’s t-tests (when data were normally distributed but not equally variable) or Mann–Whitney tests (when a non-parametric test was appropriate). Multiple t-tests (parametric or non-parametric) were followed by the False Discovery Rate (FDR) multiple comparisons adjustment. To assess the statistical significance of differences in median fluorescence intensity of CellRox Green staining, we used a Welch’s one-way ANOVA (when data were normally distributed but unequally variable) followed by the Dunnett’s T3 method for multiple comparisons adjustment. To assess the statistical significance of differences in metabolite levels between subcutaneous and metastatic tumors we used repeated measures two-way ANOVAs (samples were matched, and a parametric test was appropriate). To assess the statistical significance of melanoma cell growth over time in culture, we used the nparLD method (62), a statistical tool for non-parametric longitudinal data analysis. To assess the statistical significance of differences between multiple control clones and multiple *G6PD^mut^* clones, we performed linear mixed effects analysis or generalized linear mixed effects analysis by combining data from A375, M214, and M481. Multiple comparisons were adjusted using the FDR method. All statistical analyses were performed using Graphpad Prism9.2.0 or R 4.0.2 with the stats, fBasics, car, lme4, emmeans, and nparLD packages. All data are mean ± s.d.

No data were excluded; however, mice sometimes died during experiments, presumably due to the growth of metastatic tumors. In those instances, data that had already been collected on the mice in interim analyses were included (such as subcutaneous tumor growth measurements over time) even if it was not possible to perform the end-point analysis of metastatic disease burden due to the premature death of the mice.

## Acknowledgments

S.J.M. is a Howard Hughes Medical Institute Investigator, the Mary McDermott Cook Chair in Pediatric Genetics, the Kathryn and Gene Bishop Distinguished Chair in Pediatric Research, the director of the Hamon Laboratory for Stem Cells and Cancer, and a Cancer Prevention and Research Institute of Texas Scholar. The research was supported by the Cancer Prevention and Research Institute of Texas (RP170114 and RP180778) and by the National Institutes of Health (NIH; U01 CA228608). J.G. was supported by the Dermatology Foundation and the National Institutes of Health (T32 AR065969). We thank M. Nitcher for mouse colony management; V. Ramesh for management of the melanoma bank; the BioHPC (High Performance Computing) for data storage; and the Moody Foundation Flow Cytometry Facility.

## Footnotes

### Author contributions

A.B.A. and S.J.M. conceived the project, and designed and interpreted experiments. A.B.A. performed most of the experiments with help from V.K., J.G.G., and A.L. J.G.G. performed RNAseq experiments and helped with the interpretation of experiments. T.P.M., M.M.S., W.G. and R.J.D. participated in the design, analysis, and interpretation of isotope tracing and metabolomics experiments. V.K. and A.L. performed in vivo assays including xenografting mice and analyzing metastatic burden by bioluminescence imaging. V.K., A.T. and S.Y.K. optimized methods for CRISPR gene-targeting in patient-derived melanomas. C.Y. developed the GDH activity assay and performed the GDH experiments. T.P.M., W.G. and M.M. performed all mass spectrometric analyses. Z.Z. performed statistical analyses. A.B.A. and S.J.M. wrote the manuscript.

## Supplementary figure legends

**Supplementary Figure 1:**
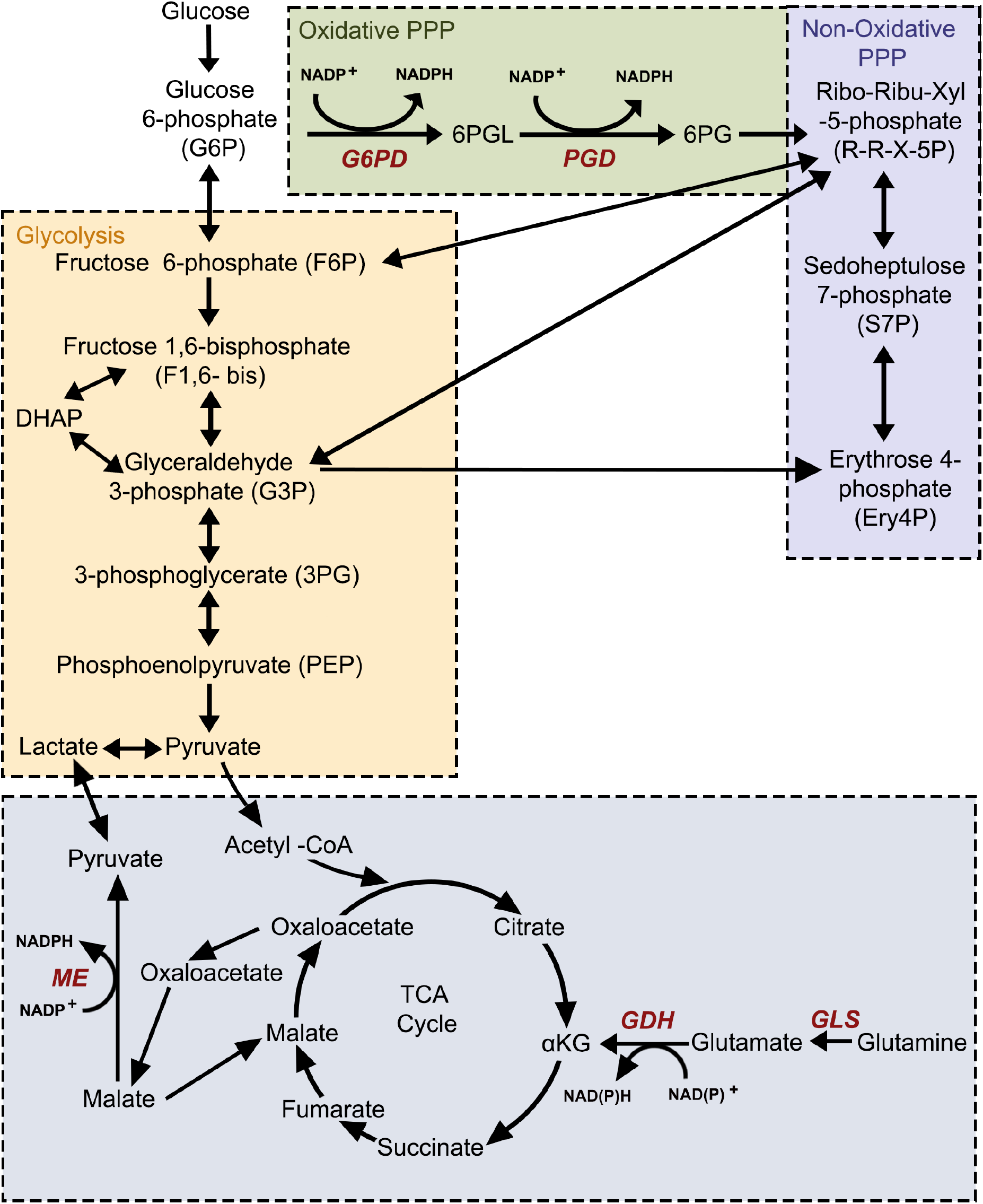
A schematic of the oxidative (green) and non-oxidative (blue) branches of the pentose phosphate pathway as well as glycolysis (yellow) and the TCA cycle (gray). The schematic includes all enzymes and metabolites mentioned in our studies.

**Supplementary Figure 2:**
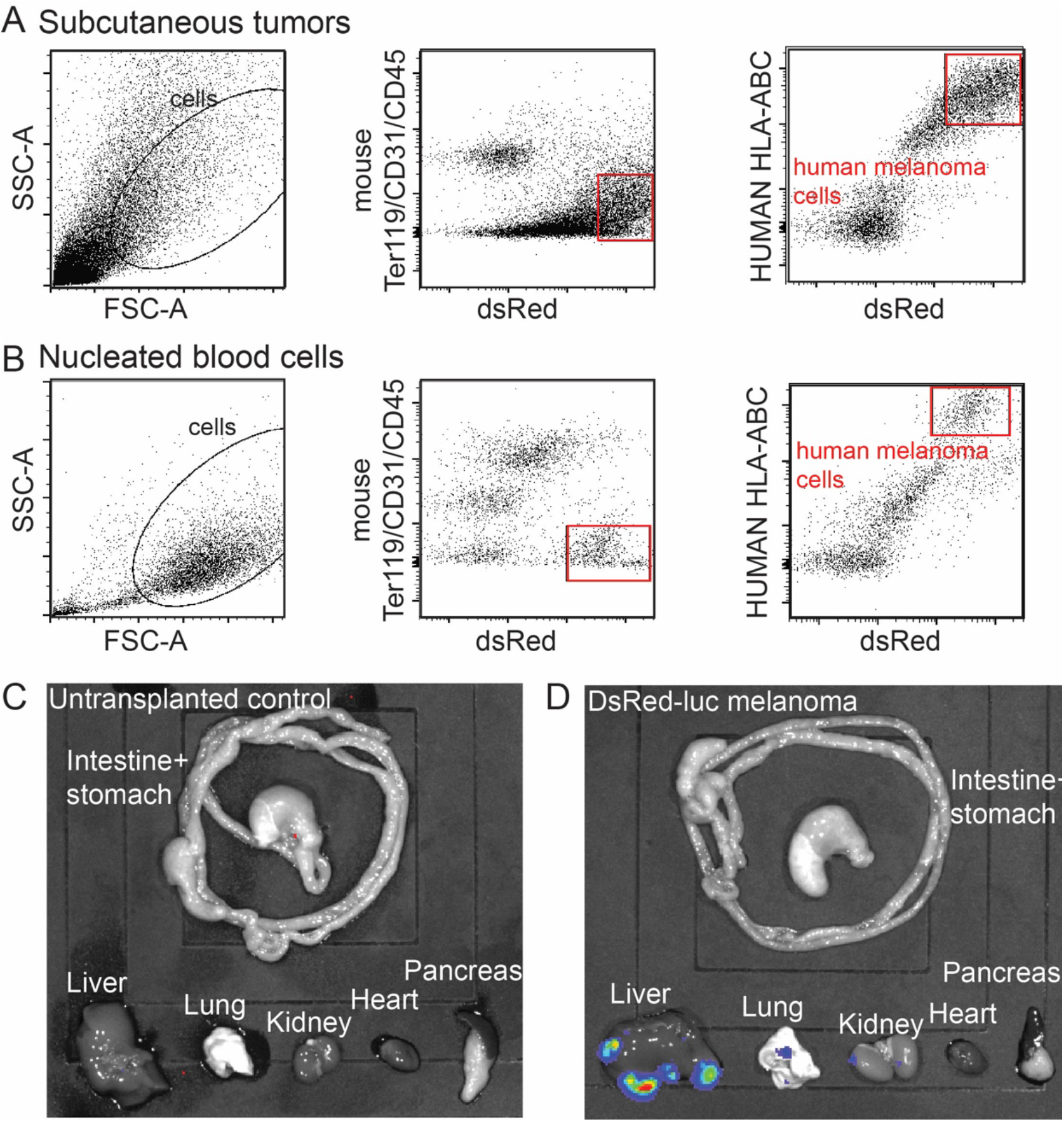
The identification of human melanoma cells by flow cytometry and assessment of metastatic disease burden by bioluminescence imaging. (A-B). Flow cytometry plots showing the gating strategies used to identify human melanoma cells in subcutaneous tumors (*A*) or nucleated blood cells (*B*) obtained from xenografted mice. Cells were gated on forward versus side scatter (FSC-A versus SSC-A) to exclude red blood cells, debris, and clusters of cells. Human melanoma cells were selected by including cells that stained positively for DsRed (stably expressed in all melanoma lines) and HLA and excluding cells that stained positively for the mouse hematopoietic and endothelial markers CD45, CD31 or Ter119. (C-D) Visceral organs were surgically removed from each mouse at the end of each experiment and imaged to identify macrometastases and micrometastases and to determine bioluminescence signal intensity. Each melanoma constitutively expressed luciferase. Shown are representative images from an untransplanted negative control mouse (*C*), illustrating background bioluminescence, and a mouse subcutaneously transplanted with dsRed-luciferase expressing melanoma cells (*D*) with metastases in the liver, lung, and kidney.

## Notes

### Competing Interest Statement

The authors have declared no competing interest.

